# A putative new SARS-CoV protein, 3a*, encoded in an ORF overlapping ORF3a

**DOI:** 10.1101/2020.05.12.088088

**Authors:** Andrew E. Firth

## Abstract

Identification of the full complement of genes in SARS-CoV-2 is a crucial step towards gaining a fuller understanding of its molecular biology. However, short and/or overlapping genes can be difficult to detect using conventional computational approaches, whereas high throughput experimental approaches – such as ribosome profiling – cannot distinguish translation of functional peptides from regulatory translation or translational noise. By studying regions showing enhanced conservation at synonymous sites in alignments of SARS-CoV and related viruses (subgenus *Sarbecovirus*), and correlating with the conserved presence of an open reading frame and plausible translation mechanism, we identified a putative new gene, ORF3a*, overlapping ORF3a in an alternative reading frame. A recently published ribosome profiling study confirmed that ORF3a* is indeed translated during infection. ORF3a* is conserved across the subgenus *Sarbecovirus*, and encodes a 40–41 amino acid predicted transmembrane protein.

## INTRODUCTION

The aetiological agent of COVID-19 is the virus SARS-CoV-2, a coronavirus in the genus *Betacoronavirus*, subgenus *Sarbecovirus*. Like other coronaviruses, SARS-CoV-2 has a positive-sense RNA genome, around 30,000 nt in size. The 5′ two thirds of the genome contain two long open reading frames (ORFs), ORF1a and ORF1b, which are translated from the viral genomic RNA (gRNA). ORFs 1a and 1b encode the polyproteins pp1a and pp1ab, where translation of pp1ab depends on a proportion of ribosomes making a programmed −1 nt ribosomal frameshift near the end of ORF1a to enter ORF1b. Polyproteins pp1a and pp1ab are proteolytically processed to produce the viral replication proteins (reviewed in Snijder et al., 2016). The 3′ third of the genome contains a number of ORFs that encode the viral structural and accessory proteins. These ORFs are translated from a nested series of subgenomic mRNAs (sgmRNAs) that are produced during the infection cycle (reviewed in Sola et al., 2015). In SARS-CoV-2, these ORFs comprise S, 3a, E, M, 6, 7a, 7b, 8, N, 9b and possibly 10 (Fig. 1a). The S, E, M and N ORFs encode respectively the virus spike, envelope, membrane and nucleocapsid proteins – key components of the virus particle that are conserved among coronaviruses. ORFs 3a, 6, 7a, 7b, 8 and 9b encode accessory proteins, also present in SARS-CoV-1, but less widely conserved across coronaviruses as a whole (reviewed in Liu et al., 2014). These 3′ ORFs each have a corresponding dedicated sgmRNA except for ORFs 7b, 9b and 10. ORFs 7b and 9b appear to be translated from the ORF7a and N sgmRNAs, respectively, via a leaky scanning mechanism (Schaecher et al., 2007; Xu et al., 2009) whereas the translation mechanism for ORF10 is unknown.

**Figure 1.**
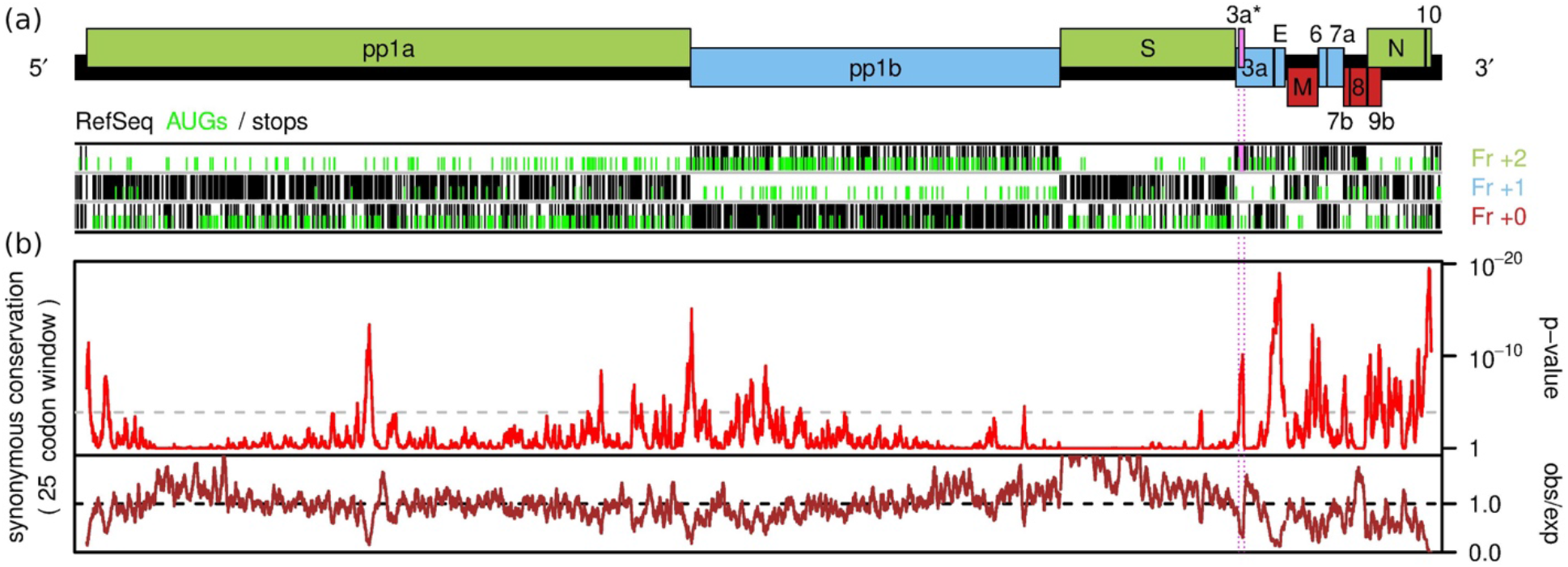
Synonymous site conservation analysis of sarbecoviruses. **(a)** Map of the SARS-CoV-2 genome (29903 nt; black rectangle). Known ORFs are overlaid in red, blue or green depending on their relative reading frames (+0, +1, +2, respectively). Below are shown the positions of AUG (green) and stop (black) codons in each of the three reading frames, as indicated, in the reference sequence NC_045512.2. The putative ORF3a* is indicated in pink. **(b)** Synonymous site conservation analysis of 54 aligned sarbecovirus sequences. The red line shows the probability that the observed conservation could occur under a null model of neutral evolution at synonymous sites, whereas the brown line depicts the ratio of the observed number of substitutions to the number expected under the null model. The horizontal dashed grey line indicates a *p* = 0.05 threshold after an approximate correction for multiple testing, namely scaling by (25-codon window size)/(length of plot in codons). Prior to analysis, the alignment was mapped to NC_045512.2 coordinates by removing alignment positions in which NC_045512.2 contained a gap character.

There is considerable variability between coronavirus genera and subgenera in the complement of 3′-encoded accessory genes (Liu et al, 2014). Even within the sarbecovirus genus, there are differences. For example, SARS-CoV-1 has an ORF3b that overlaps the 3′ region of ORF3a but is absent in SARS-CoV-2. Also, in many human-adapted SARS-CoV-1 isolates, ORF8 is split by a frame-disrupting deletion (He et al., 2004). ORF10 is apparently translated in SARS-CoV-2 (Finkel et al., 2020) but is truncated in SARS-CoV-1. Identification of the full complement of genes in SARS-CoV-2 is a crucial step towards gaining a fuller understanding of its molecular biology, and may also help guide vaccine or other antiviral strategies. This information also facilitates rational manipulation of the viral genome (e.g. for developing replicon systems or for mutagenesis studies). However, short and/or overlapping genes can be particularly difficult to identify using traditional computational approaches. On the other hand, high throughput experimental techniques such as ribosome profiling and high resolution mass spectrometry – while powerful – do not necessarily distinguish between functional proteins, regulatory translation (where it is the act of translation rather than the encoded product that is biologically relevant), and translational noise.

Comparative genomics offers a way forward: analysis of patterns of substitutions across alignments of related sequences can be used to reveal the signatures of “hidden” protein-coding genes. Analysis of synonymous substitution rates provides a particularly sensitive technique for identifying *overlapping* functional elements embedded within protein-coding genes, because such elements constrain synonymous changes which otherwise are selectively more-or-less neutral (Firth, 2014). When combined with the conserved presence of an open reading frame and the conserved presence of a plausible translation mechanism, overlapping genes may be distinguishable from overlapping non-coding elements such as functionally important RNA structures (Jagger et al., 2012; Fang et al., 2012; Loughran et al., 2012; Lulla & Firth, 2020).

## RESULTS & DISCUSSION

A full genome alignment of 54 representative sarbecoviruses (SARS-CoV-1, SARS-CoV-2 and 52 bat coronaviruses) was constructed, and synonymous site conservation was analysed with the program synplot2 (Firth, 2014) (Fig. 1b). The analysis revealed conserved features towards the 5′ end of ORF1a, in the middle of ORF1a, near the end of ORF1a and start of ORF1b, and in many parts of the 3′ region of the viral genome. Most of these conserved elements do not correspond to conserved open reading frames and likely represent functional RNA sequence elements – a case in point being the frameshift-stimulating RNA pseudoknot (Baranov et al., 2005) at the junction of ORFs 1a and 1b. However a peak in conservation within ORF3a stood out since, although short, it coincides with an overlapping alternative-frame AUG-initiated ORF – hereafter ORF3a* – positioned close to the 5′ end of ORF3a where it might be accessible via ribosomal leaky scanning (reviewed in Firth & Brierley, 2012).

Closer inspection revealed that the presence and location of the 3a* initiation and stop codons were conserved across sarbecoviruses. The 3a* AUG codon is present in all except one of the 54 sequences, where it is replaced with GUG (MG772933, a bat CoV). In all 54 sequences, there is an A at the −3 position giving a strong initiation context (Kozak, 1986). The GUG in MG772933 may also serve as an initiation site as GUG codons can be utilized for initiation – at a reduced efficiency compared to AUG – when in a strong initiation context (reviewed in Firth & Brierley, 2012). In 52 sequences there are two upstream AUGs – the ORF3a AUG (weak context; C or U at −3) and another AUG also in the ORF3a reading frame (weak context; U at −3); in the remaining two sequences the second of these AUGs is absent. The two weak-context AUGs might enable a proportion of pre-initiation scanning 43S complexes loaded on the ORF3a sgmRNA to leaky scan to the 3a* initiation codon. The 3a* ORF has length 40 codons in all 54 sequences except one, namely SARS-CoV-2, where it is 41 codons in length (Fig. 2). The SARS-CoV-2 putative 3a* protein has a molecular mass of 4.9 kDa and a pI of 10.9. The protein is also predicted to contain a transmembrane domain (Fig. 2). Curiously, the transmembrane amino acids are relative highly conserved, suggesting they may form interactions within the lipid bilayer for example membrane-disrupting or membrane-associated signalling activities.

**Figure 2.**
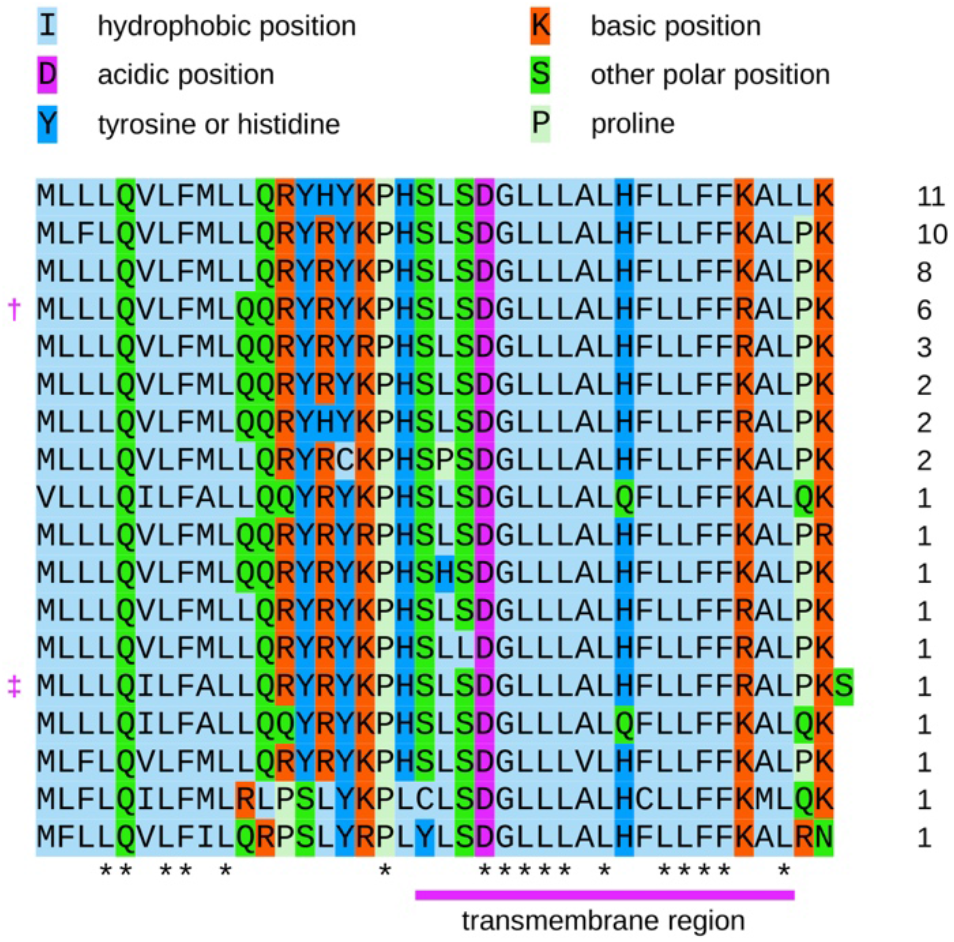
Amino acid alignment of sarbecovirus 3a* sequences. Amino acids are colour-coded according to their physicochemical properties. Asterisks indicate completely conserved columns in the alignment. The predicted transmembrane region is indicated with a pink bar below the alignment. Numbers at right indicate the number of times the particular sequence occurs among the 54 sarbecovirus sequences. † SARS-CoV-1. ‡ SARS-CoV-2. For the sequence beginning with GUG instead of AUG, the genetic decoding (i.e. valine) is shown, even though non-AUG initiation codons are normally expected to be decoded as methionine by initiator Met-tRNA.

Ribosome profiling is a high throughput sequencing technique that allows footprinting of initiating and/or elongating ribosomes at sub-codon resolution and hence the global identification of initiation sites and/or sequence regions and reading frames undergoing translation (Ingolia et al., 2009; Irigoyen et al., 2016). A recent ribosome profiling study of cells infected with SARS-COV-2 revealed 23 novel translated ORFs (Finkel et al., 2020). Ten are very short (≤ 15 codons). Seven of the remainder comprise 5′ extensions or 5′ truncations of previously known ORFs (M, 6, 7a, 7b, 9b and 10). Two are uORFs positioned on the gRNA upstream of ORF1a that may play a role in regulating ORF1a/1b expression as previously proposed for uORFs in other coronaviruses (Wu et al., 2014; Irigoyen et al., 2016). After excluding these ORFs, only four novel translated ORFs remain: ORF3a* (25457‥25582; 41 codons), another ORF overlapping ORF3a (25596‥25697; 33 codons), an ORF overlapping the S ORF (21744‥21863; 39 codons) and a truncated version of the same (21768‥21863; 31 codons) (where numbers indicate ORF coordinates in NC_045512.2). We investigated the four overlapping ORFs (yellow rectangles, Fig. 3) and found only ORF3a* to coincide with a synonymous site conservation signal. Moreover, the other novel ORFs were not conserved: in many sarbecovirus sequences they lacked the AUG codon and were interrupted by stop codons (Fig. 3).

**Figure 3.**
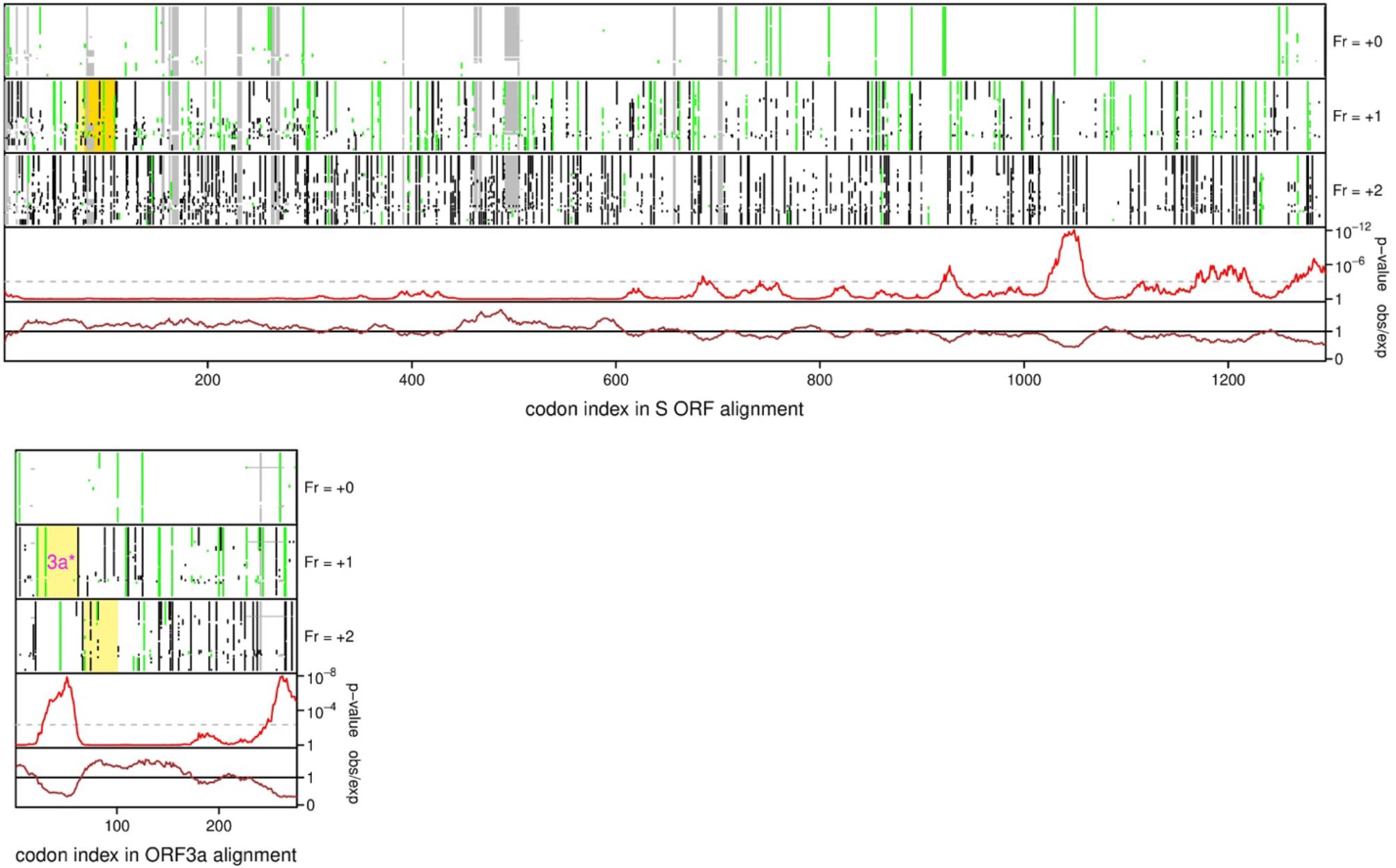
Conservation analyses of the sarbecovirus S and 3a ORFs. In each plot, the upper three panels show the positions of alignment gaps (grey), stop codons (black) and AUG codons (green) in each reading frame in each of the 54 aligned sequences. Below, is shown the analysis of conservation at synonymous sites (see Fig. 1 caption for details). Novel alternative-frame translated ORFs identified in the SARS-CoV-2 ribosome profiling study of Finkel et al. (2020) are indicated with yellow rectangles; for S, the two in-frame alternative initiation sites are indicated with different shades of yellow. ORF3a* is labelled. Note that only ORF3a* has conserved start and stop codon positions across sarbecoviruses and only ORF3a* coincides with a region of enhanced synonymous site conservation.

Although synonymous site conservation can result from overlapping non-coding or coding elements, the conserved presence and conserved positions of the ORF3a* start and stop codons suggests the latter interpretation. Moreover, the ribosome profiling study of Finkel et al. (2020) confirms that the ORF3a* is indeed translated during infection. The combination of comparative genomics showing purifying selection (which to a large extent is synonymous with functional importance) and ribosome profiling showing expession strongly suggests that 3a* is a functional protein, conserved throughout sarbecoviruses. While the known SARS-CoV-2 genes have already been studied in SARS-CoV-1 (reviewed in Liu et al., 2014), 3a* has never before been studied. Clearly additional work with SARS-CoV reverse genetics systems will be required to elucidate the 3a* protein function, but it may eventually provide a new target for vaccine or antiviral strategies. The synplot2 analysis (Fig. 1b) can also reveal other functional elements embedded within the viral protein-coding genes, which may also be worthy of experimental investigation.

During preparation of this manuscript, 3a* was independently discovered by Cagliani et al., (2020) (where it is termed 3h) who performed a similar analysis with synplot2 but using far fewer sarbecovirus sequences, and hence achieving lower statistical significance for conserved elements.

## METHODS

Sequences with 100% coverage of, and ≥70% amino acid identity to the SARS-CoV-2 pp1ab sequence were identified with NCBI tblastn (Altschul et al., 1990) on 12 January 2020 and downloaded. These cut-off thresholds corresponded precisely to genus *Sarbecovirus*. Sequences that did not cover the entire protein-coding region of the genome, and all sequences with >99% amino acid identity in pp1ab to SARS-CoV-1 except a single reference sequence NC_004718.3 were removed. This left 54 sequences (SARS-CoV-1, SARS-CoV-2 and 52 bat CoVs) with GenBank accession numbers NC_045512.2, AY274119.3, DQ022305.2, DQ071615.1, DQ084199.1, DQ084200.1, DQ412042.1, DQ412043.1, DQ648856.1, DQ648857.1, GQ153539.1, GQ153540.1, GQ153541.1, GQ153542.1, GQ153543.1, GQ153544.1, GQ153545.1, GQ153546.1, GQ153547.1, GQ153548.1, GU190215.1, JX993987.1, JX993988.1, KC881006.1, KF294457.1, KF367457.1, KF569996.1, KJ473811.1, KJ473812.1, KJ473813.1, KJ473816.1, KP886808.1, KP886809.1, KT444582.1, KY352407.1, KY417142.1, KY417143.1, KY417145.1, KY417146.1, KY417147.1, KY417148.1, KY417149.1, KY417150.1, KY417151.1, KY417152.1, KY770858.1, KY770859.1, KY770860.1, MG772933.1, MG772934.1, MK211374.1, MK211376.1, MK211377.1 and MK211378.1.

Synonymous site conservation was analyzed with synplot2 (Firth, 2014). For the full-genome analyses, codon-respecting alignments were produced using a procedure described previously (Firth, 2014). In brief, each individual genome sequence was aligned to the SARS-CoV-2 reference sequence (GenBank accession number NC_045512.2) using code2aln version 1.2 (Stocsits et al., 2005). Genomes were then mapped to NC_045512.2 coordinates by removing alignment positions that contained a gap character in the reference sequence, and these pairwise alignments were combined to give the multiple sequence alignment. To assess conservation at synonymous sites, the known virus coding regions were extracted from the alignment (with codons selected from the longer ORF in each overlap region), concatenated in-frame, and the alignment analyzed with synplot2 using a 25-codon sliding window. Conservation statistics were then mapped back to NC_045512.2 coordinates for plotting (Fig. 1).

For the ORF S and 3a analyses (Fig. 3), the ORF S and 3a regions were extracted from all 54 sarbecovirus sequences, translated to amino acids, aligned using MUSCLE (Edgar, 2004), and the amino acid alignments were used to guide codon-respecting nucleotide sequence alignments (EMBOSS tranalign; Rice et al., 2000). These alignments were analyzed with synplot2, again using a 25-codon window size. In contrast to the full-genome alignments, all alignment gaps were retained instead of mapping to NC_045512.2 coordinates.

Molecular mass and isoelectric point were calculated with EMBOSS pepstats (Rice et al., 2000). Transmembrane domains were predicted with Phobius (Käll et al., 2007).

## ACKNOWLEDGEMENTS

This work was supported by Wellcome Trust grant [106207] and European Research Council grant [646891] to A.E.F. I thank Valeria Lulla, Hazel Stewart, Betty Chung, James Edgar, Ian Brierley, Andrew Davidson, David Matthews and Nina Lukhovitskaya for useful discussions.

## REFERENCES

Altschul SF, Gish W, Miller W, Myers EW, Lipman DJ (1990) Basic local alignment search tool. J Mol Biol 215:403–410. PMID 2231712.

Baranov PV, Henderson CM, Anderson CB, Gesteland RF, Atkins JF, Howard MT (2005) Programmed ribosomal frameshifting in decoding the SARS-CoV genome. Virology 332:498–510. PMID 15680415.

Cagliani R, Forni D, Clerici M, Sironi M (2020) Coding potential and sequence conservation of SARS-CoV-2 and related animal viruses. Infect Genet Evol [Epub ahead of print]. PMID 32387562

Edgar RC (2004) MUSCLE: a multiple sequence alignment method with reduced time and space complexity. BMC Bioinformatics 5:113. PMID 15318951

Fang Y, Treffers EE, Li Y, Tas A, Sun Z, van der Meer Y, de Ru AH, van Veelen PA, Atkins JF, Snijder EJ, Firth AE (2012) Efficient −2 frameshifting by mammalian ribosomes to synthesize an additional arterivirus protein. Proc Natl Acad Sci U S A 109:E2920–8. PMID 23043113.

Finkel Y, Mizrahi O, Nachshon A, Weingarten-Gabbay S, Yahalom-Ronen Y, Tamir H, Achdout H, Melamed S, Weiss S, Isrealy T, Paran N, Schwartz M, Stern-Ginossar N (2020) The coding capacity of SARS-CoV-2. bioRxiv 2020.05.07.082909; doi: https://doi.org/10.1101/2020.05.07.082909

Firth AE, Brierley I (2012) Non-canonical translation in RNA viruses. J Gen Virol 93:1385–1409. PMID 22535777.

Firth AE (2014) Mapping overlapping functional elements embedded within the protein-coding regions of RNA viruses. Nucleic Acids Res 42:12425–12439. PMID 25326325.

He J et al., Chinese SARS Molecular Epidemiology Consortium (2004) Molecular evolution of the SARS coronavirus during the course of the SARS epidemic in China. Science 303:1666–1669. PMID 14752165.

Ingolia NT, Ghaemmaghami S, Newman JR, Weissman JS (2009) Genome-wide analysis in vivo of translation with nucleotide resolution using ribosome profiling. Science 324:218–223. PMID 19213877.

Irigoyen N, Firth AE, Jones JD, Chung BY, Siddell SG, Brierley I (2016) High-resolution analysis of coronavirus gene expression by RNA sequencing and ribosome profiling. PLoS Pathog 12:e1005473. PMID 26919232.

Jagger BW, Wise HM, Kash JC, Walters KA, Wills NM, Xiao YL, Dunfee RL, Schwartzman LM, Ozinsky A, Bell GL, Dalton RM, Lo A, Efstathiou S, Atkins JF, Firth AE, Taubenberger JK, Digard P (2012) An overlapping protein-coding region in influenza A virus segment 3 modulates the host response. Science 337:199–204. PMID 22745253.

Käll L, Krogh A, Sonnhammer EL (2007) Advantages of combined transmembrane topology and signal peptide prediction – the Phobius web server. Nucleic Acids Res 35:W429–432. PMID 17483518.

Kozak M (1986) Point mutations define a sequence flanking the AUG initiator codon that modulates translation by eukaryotic ribosomes. Cell 44:283–292. PMID 3943125.

Liu DX, Fung TS, Chong KK, Shukla A, Hilgenfeld R (2014) Accessory proteins of SARS-CoV and other coronaviruses. Antiviral Res 109:97–109. PMID 24995382.

Loughran G, Firth AE, Atkins JF (2011) Ribosomal frameshifting into an overlapping gene in the 2B-encoding region of the cardiovirus genome. Proc Natl Acad Sci U S A 108:E1111–9. PMID 22025686.

Lulla V, Firth AE (2019) A hidden gene in astroviruses encodes a cell-permeabilizing protein involved in virus release. bioRxiv 661579; doi: https://doi.org/10.1101/661579

Rice P, Longden I, Bleasby A (2000) EMBOSS: the European Molecular Biology Open Software Suite. Trends Genet 16:276–277. PMID 10827456.

Schaecher SR, Mackenzie JM, Pekosz A (2007) The ORF7b protein of severe acute respiratory syndrome coronavirus (SARS-CoV) is expressed in virus-infected cells and incorporated into SARS-CoV particles. J Virol 81:718–731. PMID 17079322.

Snijder EJ, Decroly E, Ziebuhr J (2016) The nonstructural proteins directing coronavirus RNA synthesis and processing. Adv Virus Res 96:59–126. PMID 27712628.

Sola I, Almazán F, Zúñiga S, Enjuanes L (2015) Continuous and discontinuous RNA synthesis in coronaviruses. Annu Rev Virol 2:265–288. PMID 26958916.

Stocsits RR, Hofacker IL, Fried C, Stadler PF (2005) Multiple sequence alignments of partially coding nucleic acid sequences. BMC Bioinformatics 6:160. PMID 15985156.

Wu HY, Guan BJ, Su YP, Fan YH, Brian DA (2014) Reselection of a genomic upstream open reading frame in mouse hepatitis coronavirus 5'-untranslated-region mutants. J Virol 88:846–858. PMID 24173235.

Xu K, Zheng BJ, Zeng R, Lu W, Lin YP, Xue L, Li L, Yang LL, Xu C, Dai J, Wang F, Li Q, Dong QX, Yang RF, Wu JR, Sun B (2009) Severe acute respiratory syndrome coronavirus accessory protein 9b is a virion-associated protein. Virology 388:279–285. PMID 19394665.

